# Conformational dynamics of FERM-mediated autoinhibition in Pyk2 tyrosine kinase

**DOI:** 10.1101/681932

**Authors:** Hanna S. Loving, Eric S. Underbakke

## Abstract

Pyk2 is a non-receptor tyrosine kinase that evolved from gene duplication of focal adhesion kinase (FAK) and subsequent functional specialization in the brain and hemopoietic cells. Pyk2 shares a domain organization with FAK, with an N-terminal regulatory FERM domain adjoining the kinase domain. FAK regulation involves integrin-mediated membrane clustering to relieve autoinhibitory interactions between FERM and kinase domains. Pyk2 regulation remains cryptic, involving Ca^2+^ influx and protein scaffolding. While the mechanism of the FAK FERM domain in autoinhibition is well-established, the regulatory role of the Pyk2 FERM is ambiguous. We probed the mechanisms of FERM-mediated autoinhibition of Pyk2 using hydrogen/deuterium exchange mass spectrometry (HDX-MS) and kinase activity profiling. The results reveal FERM-kinase interfaces responsible for autoinhibition. Pyk2 autoinhibition impacts activation loop conformation. In addition, the autoinhibitory FERM-kinase interface exhibits allosteric linkage with the FERM basic patch conserved in both FAK and Pyk2.

**Table of Contents graphic:** 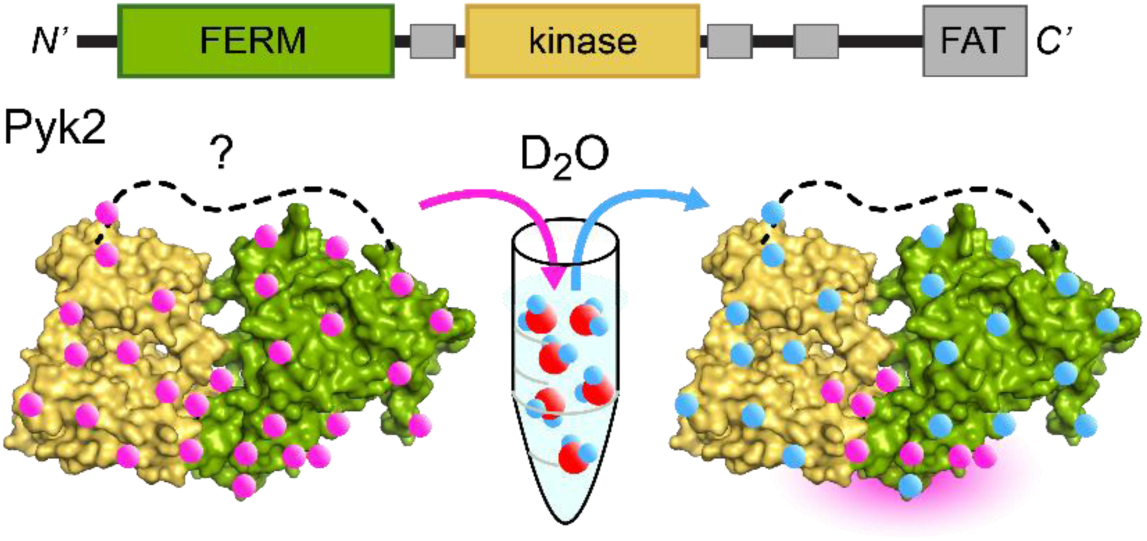

## Introduction

Proline-rich tyrosine kinase 2 (Pyk2, UniprotKB Q14289), also known as CAKβ, FAK2, or RAFTK, is a non-receptor tyrosine kinase and paralogue of focal adhesion kinase (FAK).^*1*^ FAK is expressed ubiquitously, while Pyk2 is expressed in a limited number of cell types, primarily central nervous system (CNS) tissues and hematopoietic cells.^*1-4*^ Pyk2 appears to have arisen in early chordates from a gene duplication of FAK, sharing 45% sequence identity (65% similarity).^*5*^ As such, Pyk2 retains functional overlap with FAK and can compensate for some FAK functions in knockout models.^*6-8*^ Nevertheless, Pyk2 function has specialized, adopting unique regulatory mechanisms and signaling roles.^*9*^ Whereas FAK is localized primarily at focal adhesions, Pyk2 exhibits diffuse cytosolic localization.^*6, 10, 11*^ FAK plays central signaling roles in cell adhesion, migration, proliferation, and embryonic development.^*12, 13*^ In contrast, Pyk2 is enriched in the post-synaptic density of neurons and serves to tune synaptic plasticity and regulate neuronal differentiation and migration in the CNS.^*14-20*^ Pyk2 signaling is also involved in bone remodeling and leukocyte motility.^*3, 21*^ Canonical FAK activation involves recruitment to membrane focal adhesions via interactions with paxillin, talin, and vinculin at clustered integrins.^*22-24*^ Strikingly, Pyk2 is activated by stimuli that elicit intracellular Ca^2+^ flux.^*1, 14, 16, 25*^ The marked differences in activation mechanisms between FAK and Pyk2 (i.e., integrin-clustering vs. Ca^2+^ flux) illustrate a common theme in cell signaling: repurposing of common domain components to respond to diverse stimuli. Understanding the molecular details of the divergent regulation of FAK and Pyk2 can illuminate how signaling diversity can arise through gene duplication and mechanistic shifts.

Pyk2 and FAK both exhibit a kinase domain flanked by a FERM (band 4.1/ezrin/radixin/moesin) domain at the N-terminus and a FAT (focal adhesion targeting) domain at the C-terminus (Figure 1). The FAT domain is tethered to the kinase via a putatively unstructured region containing several proline-rich regions. The multi-domain architecture is punctuated by binding motifs, allowing Pyk2 and FAK to serve signaling roles as both kinase and scaffold.^*22, 26-28*^

**Figure 1.**
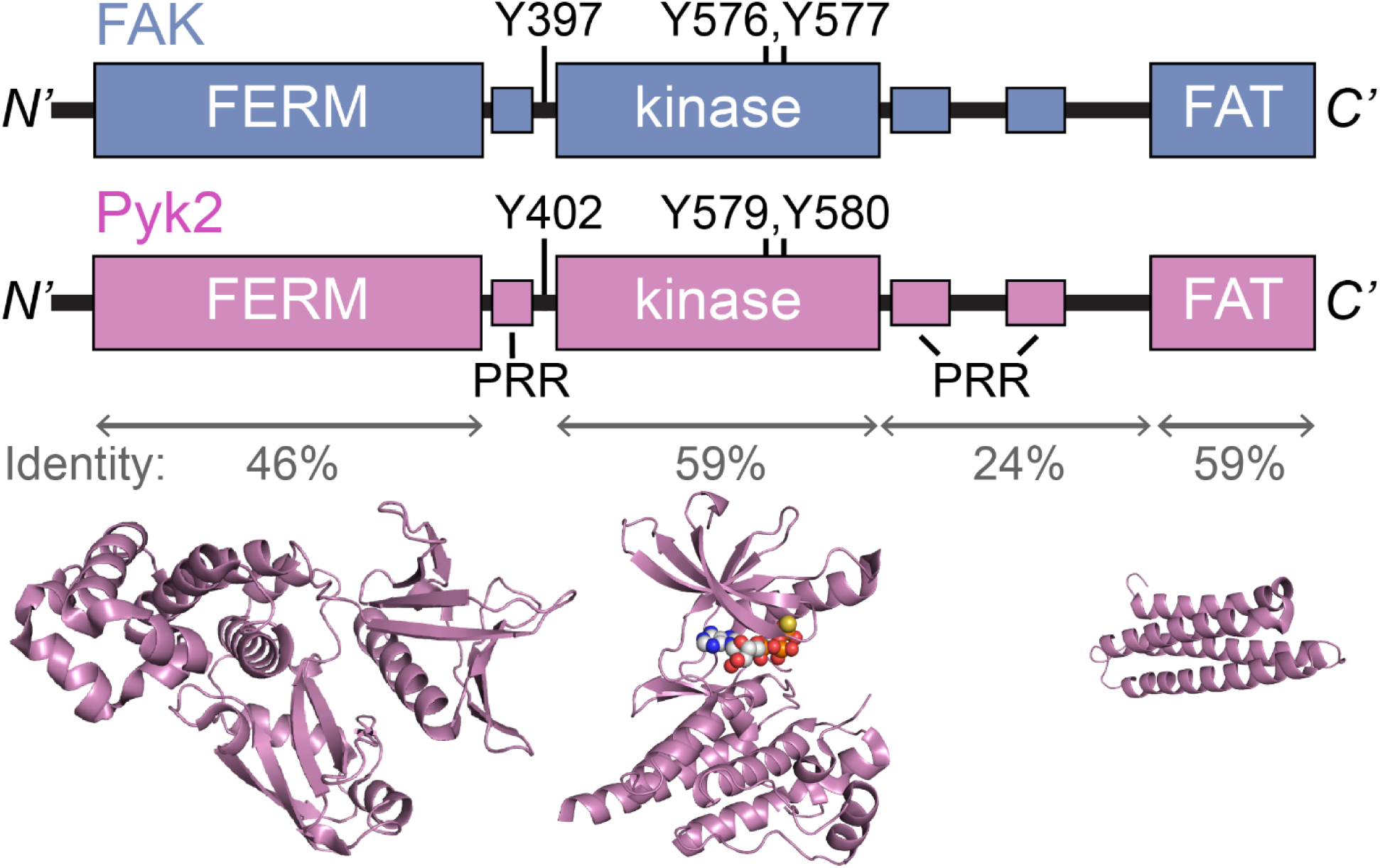
(A) Domain organization of Pyk2 and FAK. Pairwise identity of the *Homo sapiens* sequences, phosphorylated tyrosine residues, and proline-rich regions (PRR) are annotated. (B) Structural models of the Pyk2 domains are depicted from the following PDB ID: FERM, 4eku; kinase 3fzp; FAT, 3gm2.^*29, 30*^

The mechanisms of FAK activation are not fully understood, but integrin-mediated clustering and membrane phosphoinositide interactions appear to relieve autoinhibitory interactions between the FERM and kinase domains.^*24, 31-34*^ The dimerization and clustering of de-repressed FAK kinase domains allows for trans-autophosphorylation of residue Y397 in the FERM—kinase linker. The FAK linker phosphotyrosine serves as a docking site for Src kinase, leading to mutual phosphorylation of FAK and Src kinase domain activation loops (FAK residues Y576, Y577). Ultimately, FAK clustering at the membrane of focal adhesions is responsible initiating the FAK signaling cascade.

The initial steps of Pyk2 activation remain unclear. Ca^2+^-induced activation may involve dimerization via Ca^2+^/calmodulin-binding^*35, 36*^ or a Ca^2+^-dependent multimerization via docking to the soluble scaffold protein PSD-95.^*16*^ Nevertheless, downstream steps parallel FAK. Clustered Pyk2 autophosphorylates in trans at a FERM—kinase linker tyrosine (Y402), followed by Src-binding and reciprocal activation loop phosphorylation for full activation.^*37*^

Clarifying the mechanism of Pyk2 autoinhibition is a prerequisite to understanding the activation mechanism. FAK autoinhibition is relatively well-defined by the x-ray crystallography-derived structure of an avian FAK construct encompassing the FERM-kinase domains (residues 31-686).^*34*^ The FAK FERM-kinase structure revealed interactions between the kinase C-lobe and FERM F2 subdomain stabilizing a closed conformation. The closed conformation occludes the active site and sequesters the initial site of phosphorylation (Y397) far from the active site.

The autoinhibitory role of the Pyk2 FERM domain is not yet resolved.^*9*^ To date, no structure of the autoinhibited Pyk2 has been reported. Cell-based studies establish a clear regulatory role for the Pyk2 FERM.^*11, 38-41*^ Point mutations in the Pyk2 FERM domain can inhibit the activity of the kinase.^*39*^ Expression of an autonomous Pyk2 FERM domain truncation inhibits the basal autophosphorylation of full-length Pyk2,^*39, 40, 42*^ an observation echoed in FAK.^*33*^ However, in some cases, Pyk2 FERM notably deviates from FAK FERM activity. Immunoprecipitation established that FAK FERM interacts with full-length FAK and the isolated FAK kinase domain, yet Pyk2 FERM exhibits negligible association with the dissociated Pyk2 kinase.^*38, 40*^ Intriguingly, FERM domain deletion strongly increases FAK autophosphorylation,^*33*^ yet FERM deletion in Pyk2 did not impact basal autophosphorylation in cells.^*41*^

To resolve the mechanism of Pyk2 autoinhibition, we probed functional interactions between FERM and kinase domains. Our approach integrates hydrogen/deuterium exchange mass spectrometry (HDX-MS) and site-directed mutagenesis to reveal inhibitory interfaces between the Pyk2 FERM and kinase domains. The investigation also informs on activation loop dynamics and allosteric coupling in the autoinhibited conformation.

## MATERIALS AND METHODS

### Plasmid Constructs

Plasmids used in the study were derived from the following sources: *Homo sapiens* Pyk2 (PTK2B [1-1010]) was acquired from the Harvard PlasmID Repository, HsCD00022350. The *H. sapiens* Pyk2 kinase domain expression vector PTK2BA encoding H_6_-TEV-kinase [414-692] was a gift from Nicola Burgess-Brown (Addgene # 42401). Cloning vector pET-H6-SUMO-TEV-LIC (1S) was a gift from Scott Gradia (Addgene # 29659). The vector encoding *Yersinia pestis* YopH phosphatase [1-468] acquired from the Harvard PlasmID Repository, YpCD00017249. The *Saccharomyces cerevisiae* SUMO protease H_6_-Ulp1 [residues 423-621] expression vector pHYRS52 was a gift from Hideo Iwai (Addgene # 31122).

Expression vectors for Pyk2 domain truncations were generated using Gibson assembly.^*43*^ Pyk2 FERM-kinase [residues 20-692] and FERM [residues 20-357] were assembled in-frame with the His_6_-SUMO solubility tag of pET-H6-SUMO-TEV-LIC (1S) to generate pHL001 and pHL002, respectively. Pyk2 FERM-kinase mutants were generated using the QuikChange (Agilent Genomics) site-directed mutagenesis strategy. All constructs were confirmed by DNA sequencing.

### Protein Purification

Expression plasmids pHL001 and pHL002 encoding H_6_-SUMO-FERM-kinase and H_6_-SUMO-FERM, respectively, were transformed into Tuner(DE3) pLysS (Novagen) cells. Plasmid PTK2BA encoding H_6_-TEV-kinase was transformed into BL21(DE3). Starter cultures in LB media supplemented with 50 ug/mL kanamycin were grown overnight at 37 °C. Four 1 L expression cultures of LB supplemented with 50 ug/mL kanamycin were inoculated (1:100) with starter culture. Expression cultures were grown at 37 °C with shaking to OD_600nm_ of 0.35, at which point the temperature was lowered to 18 °C. At OD_600nm_ of 0.5, the cells were induced with 1 mM IPTG and incubated overnight at 18 °C with shaking. Cells were harvested after 18 hr by centrifugation at 3,700 ×g for 20 mins. Cell pellets were stored at −80 °C until purification.

Harvested cells were thawed on ice with lysis buffer (150 mM NaCl, 50 mM HEPES, 15 mM imidazole, 1 mM PMSF, 20 mM EDTA, 5% glycerol, 5 mM β-mercaptoethanol (βME) pH 8) supplemented with protease inhibitor cocktail II (Research Products International). Cells were lysed by six passages through an Emulsiflex-C3 homogenizer at 15,000 psi. The lysate was cleared of cell debris by centrifugation at 20,000 ×g for 45 mins at 4 °C. Cleared lysate was passed through a HisPur^™^ Ni-NTA Superflow agarose column (2 mL resin, Thermo Scientific) pre-equilibrated in binding buffer [150 mM NaCl, 50 mM HEPES, 20 mM imidazole, 5% glycerol, 5 mM βME pH 8]. The Ni-NTA column was washed with fifteen column volumes of wash buffer (250 mM NaCl, 50 mM HEPES, 5% glycerol, 20 mM imidazole, 5 mM βME pH 8). His-tagged Pyk2 constructs were eluted with high imidazole buffer (150 mM NaCl, 50 mM HEPES, 150 mM imidazole, 5% glycerol, 5 mM β-ME pH 8). Protein-containing elution fractions were dialyzed for 3 hr with tag-specific protease, H_6_-Ulp1 for SUMO-tagged constructs or H_6_-TEV protease for H_6_-TEV-tagged kinase domain. Purification tags and proteases were removed by a subtractive second pass through the Ni-NTA column, re-equilibrated in binding buffer. Pyk2 constructs were further purified by gel filtration chromatography (GFC) using a Superdex 200 10/300 (GE Healthcare) column pre-equilibrated in GFC buffer (150 mM NaCl, 50 mM HEPES, 10% glycerol, 1 mM DTT pH 7.4). The purified proteins were concentrated using a Spin-X concentrator of 10K MWCO PES (Corning). Concentrated proteins were aliquoted, snap frozen in liquid N_2_, and stored at −80 °C.

### Kinase Assays

Kinase reactions were performed using a final concentration of 1 µM of each Pyk2 construct in buffer consisting of 150 mM NaCl, 50 mM HEPES, 8 mM MgCl_2_, 2 mM TCEP, 1 mM Na_3_VO_4_, 10% glycerol pH 7.4. Prior to the kinase reactions, the Pyk2 constructs were dephosphorylated with 0.1 µM H_6_-YopH on ice for 30 min. Kinase reactions were initiated by addition of 1.6 mM ATP (final concentration). Reactions were quenched with SDS-PAGE loading dye followed by heat denaturation at 60 °C for 10 min. Phosphotyrosine production was assessed via anti-phosphotyrosine Western blotting. Briefly, autophosphorylation reactions were separated on SDS-PAGE gels and transferred to 0.2 µm nitrocellulose membranes at 25 V for 7 mins using Trans-Blot Turbo Transfer system (Bio-Rad). Membranes were blocked with BSA and incubated with primary mouse anti-phosphotyrosine antibody (PY20, Thermo Scientific), followed by goat-anti-mouse HRP-conjugated secondary antibody (GOXMO HRP, Novex). Site-specific anti-phosphotyrosine blotting was performed with anti-phospho-PTK2B pTyr^402^ (SAB4300173, Sigma) or anti-phospho-PYK2 pTyr^579^, pTyr^580^ (44-636G, Invitrogen) primary antibodies with goat-anti-rabbit HRP-conjugated secondary antibody (GOXRB HRP, Novex). Blots were imaged with enhance chemiluminescent substrate (Pierce).

Enzyme-coupled, continuous kinase assays were performed as previously described.^*34, 44*^ Purified Pyk2 variants were dephosphorylated by pre-incubation with 0.1 µM YopH for 30 min at 25 °C. Kinase activity was initiated by adding 0.5 mM ATP to reactions containing 1 µM Pyk2 FERM-kinase variant, Glu:Tyr (4:1) polypeptide substrate (Sigma), 0.28 mM NADH, 100 µM phosphoenolpyruvate, 100 units/mL lactate dehydrogenase and 80 units/mL pyruvate kinase (Sigma) in buffer composed of 50 mM HEPES pH 7.4, 150 mM NaCl, 8 mM MgCl_2_, 5% glycerol at 30 °C. NADH consumption was monitored at 340 nm with a Cary 60 UV-Vis spectrophotometer (Agilent).

### HDX-MS mapping

Deuterium exchange reactions (27 uL) were initiated by diluting 50 pmol of Pyk2 constructs encompassing FERM, kinase, or FERM-kinase into D_2_O buffer composed of a final concentration of 150 mM KCl, 50 mM HEPES, 2 mM DTT, pD 7.4, 90% D_2_O. Exchange reactions were incubated at 24 °C. At various time points (10 sec, 45 sec, 3 min, 10 min, 30 min, 1 hr, 3 hr, 18 hr), labeling reactions were quenched to pH 2.5 by addition of 4 M guanidium chloride, 85 mM potassium phosphate (final concentrations) and snap frozen in liquid N_2_. Quenched exchange reactions were stored at −80 °C until LC-MS analysis. All exchange reactions were performed as three technical replicates.

For HDX-MS analysis, frozen exchange samples were quickly thawed and immediately injected (50 uL) into a temperature-controlled ACQUITY UPLC M-class HDX platform coupled in-line to an ESI-Q-TOF Synapt G2-Si (Waters) Mobile phases consisted of solvents A (HPLC-grade aqueous 0.1% formic acid) and B (HPLC-grade acetonitrile, 0.1% formic acid). Samples were digested by flow through an in-line Enzymate BEH-immobilized pepsin column (5 µm particle, 300 Å pore, 2.1 × 30 mm; Waters) at 25 °C with a flow rate of 50 uL/min. Peptide products were accumulated on an Acquity UPLC BEH C18 VanGuard trap column (1.7 µm, 130 Å, 2.1 × 5 mm; Waters) and desalted with 100% solvent A for 1.5 min at a flow rate of 120 uL/min. Peptides were subsequently resolved on an Acquity UPLC BEH C18 analytical column (1.7 µm, 130 Å, 1 × 100 mm; Waters) using a 7 min linear gradient of 6% to 35% solvent B. MS data was collected in positive ion, MS^E^ continuum, resolution mode with an m/z range of 50 – 2000. Ion mobility was used to further resolve peptides in the gas phase. Peptides were fragmented by argon gas collision for data-independent acquisition.

Pepsin-derived peptides were identified by fragmentation data using ProteinLynx Global Server (PLGS ver. 3.0.3, Waters). The HDX data were analyzed with DynamX ver. 3.0 software (Waters). Peptides identified by PLGS were manually curated after establishing thresholds of identifications in 2 of 3 unlabeled samples and 0.2 fragments per residue. Relative deuterium incorporation for each peptide were calculated by DynamX after manual inspection of the spectra of each isotope envelope. Differences in deuterium incorporation between HDX treatments were assessed for representative time-points approximating the actively exchanging regime of the time course. Statistical significance was determined using a two-tailed, unpaired *t* test. HDX-MS summary statistics are reported in Supporting Information Table 1 in accordance with community-based recommendations.^*45*^

## RESULTS AND DISCUSSION

Given the well-established regulatory role of the FERM domain in FAK^*33, 34*^, we tested whether the FERM domain is responsible for kinase autoinhibition in Pyk2. Accordingly, we compared autophosphorylation rates of purified Pyk2 truncations encompassing the FERM-kinase (residues 20-692) and the isolated kinase (residues 414-692). Both Pyk2 FERM-kinase and kinase constructs exhibited background tyrosine phosphorylation as purified from *E. coli*, likely due to autophosphorylation induced by the high intracellular concentrations during over-expression.^*37*^ Pyk2 constructs were dephosphorylated prior to kinase assays by pre-treating with YopH tyrosine phosphatase. Kinase activity time courses revealed that deletion of the FERM domain allows for a dramatically increased autophosphorylation of the Pyk2 kinase domain (Figure 2a). The FERM-kinase construct exhibits a considerably slower basal autophosphorylation rate. Western blotting with site-specific antibodies for Pyk2 phosphotyrosines confirmed that FERM-kinase autophosphorylation is predominantly localized to linker residue Y402 (Figure S1).

**Figure 2.**
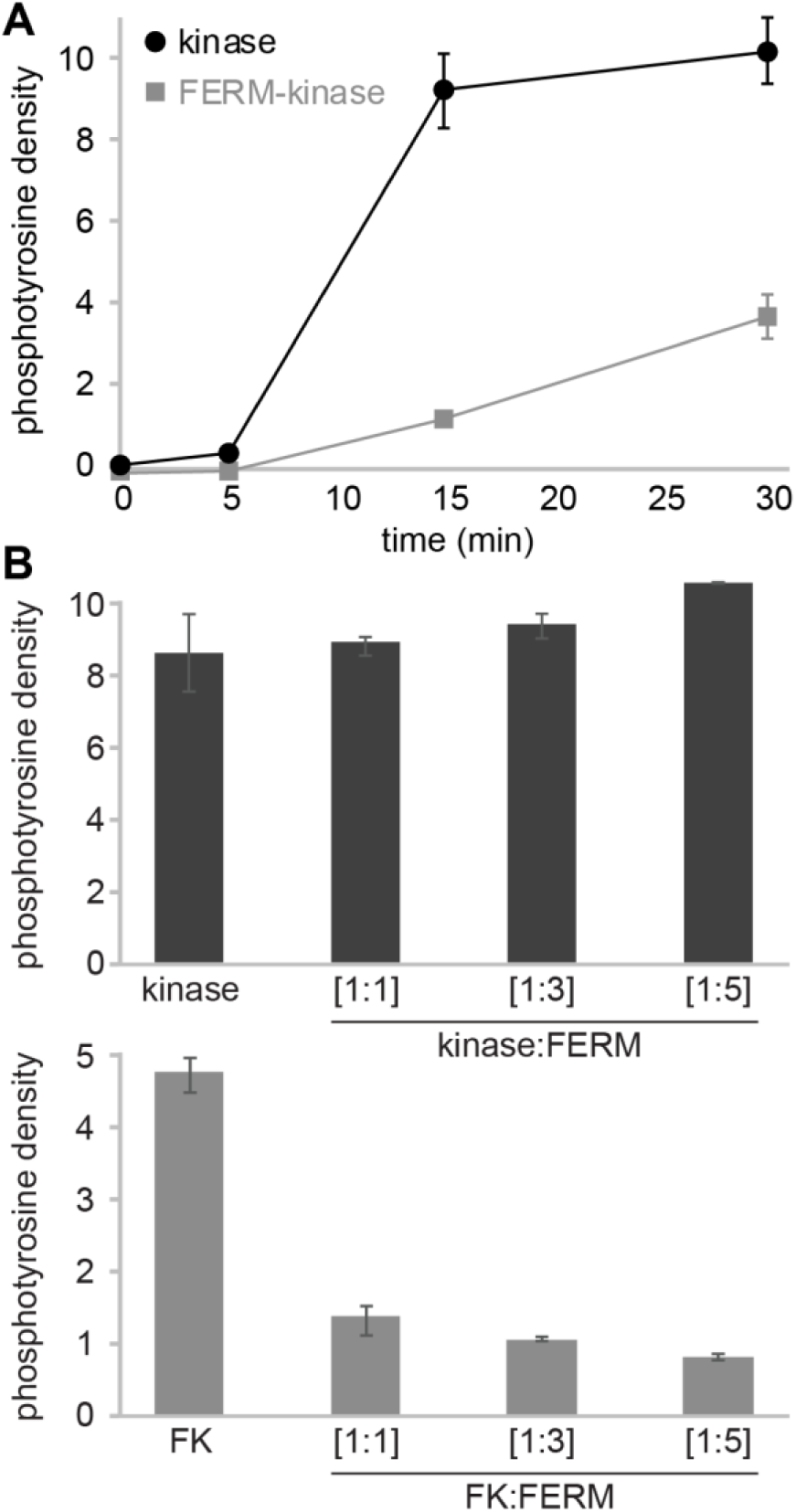
Pyk2 FERM domain inhibits kinase autophosphorylation *in vitro*. (A) Kinase activity of Pyk2 kinase and FERM-kinase constructs (1 µM) was initiated by addition of ATP. Autophosphorylation was detected by Western blotting using anti-phosphotyrosine antibody [PY20] and quantified by densitometry. (B) Relative activity of autonomous Pyk2 kinase or FERM kinase (FK) pre-incubated with buffer or exogenous FERM (1 µM, 3 µM, or 5 µM). Autophosphorylation was assessed after 30 min at 25 °C. Error bars represent standard deviation of three independent reactions.

Next, we tested whether FERM-mediated inhibition of the Pyk2 kinase requires the native intramolecular tethering. We tested the autophosphorylation activity of the isolated Pyk2 kinase incubated with autonomous FERM domain (residues 20-357) added in trans at various stoichiometries (Figure 2b). Addition of autonomous FERM domain up to five-fold molar excess does not significantly inhibit Pyk2 kinase, suggesting that the regulatory interface between FERM and kinase requires the native linker for impactful affinity. In contrast, previous studies established that autonomous FAK FERM can inhibit the FAK kinase in trans.^*33*^ Furthermore, addition of excess FAK FERM domain also suppresses basal activity of the autoinhibited, full-length FAK.^*33*^ To test whether Pyk2 FERM-kinase is sensitive to exogenous addition of FERM domain, we tested FERM-kinase autophosphorylation with various stoichiometries of added FERM domain. Indeed, free FERM domain accentuates FERM-mediated kinase inhibition in the Pyk2 FERM-kinase construct (Figure 2b).

We sought to map inter-domain interfaces responsible for FERM-mediated kinase inhibition in Pyk2. HDX-MS was used to compare the structural dynamics of Pyk2 FERM-kinase with the autonomous FERM or kinase domains. HDX-MS is a useful technique for studying protein folding, protein–protein interactions, allostery, and conformational dynamics^*46-50*^. HDX-MS measures the rates of backbone amide proton exchange to report on local chemical environments. H/D exchange rates are influenced by secondary structure, hydrogen bonding, dynamics, and solvent exposure. Increased rates of deuterium exchange result from disruptions in backbone hydrogen bonding, increased dynamics, or greater solvent accessibility. Decreased rates of exchange suggest local stabilization (e.g., formation of secondary structure and hydrogen bonding) or lower solvent accessibility. As such, exchange rate perturbations are valuable indicators of local protein dynamics, interfaces, and allostery.

We assessed the structural perturbations induced by the Pyk2 autoinhibitory FERM-kinase conformation by comparing H/D exchange rates with the free FERM and kinase domains. Changes in exchange rates were color-coded and mapped to the Pyk2 primary sequence (Figure 3a) and structural models of isolated FERM and kinase domains (Figure 3b). HDX-MS revealed several regions in both FERM and kinase impacted by the autoinhibited conformation.

**Figure 3.**
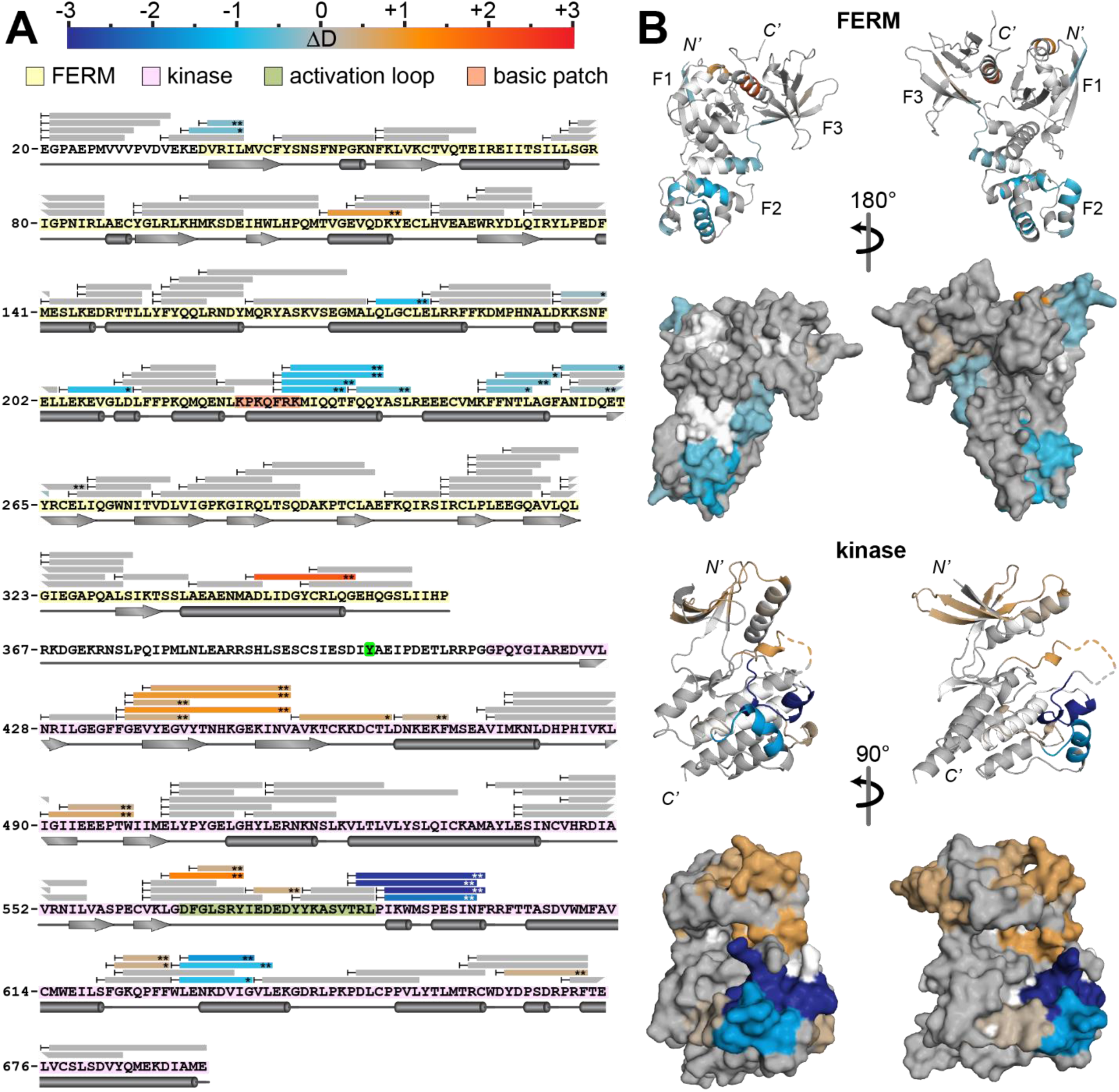
Comparison of HDX-MS exchange kinetics of Pyk2 FERM-kinase with autonomous FERM and kinase domains. (A) Peptide coverage is represented as bars mapped to primary sequence. Exchange rate perturbations are reported as the average difference in deuterium incorporation (ΔD) at time points approximating the midpoint of exchange. Peptides exhibiting significant exchange rate perturbations are color-coded according to the scale bar (top). Slower exchange in the FERM-kinase relative to autonomous FERM or kinase is reported in blue shades, while more rapid exchange in FERM-kinase is in red shades. Significance was assessed with a two-tailed, unpaired Student’s *t* test (* *p* < 0.005, ** *p* < 0.001). Peptides exhibiting negligible differences are colored gray. (B) Color-coded exchange rate perturbations were mapped to reported structures of the isolated Pyk2 FERM (PDB 4eku) and kinase (PDB 3cc6) domains. N’-, C’-termini and FERM subdomains (F1, F2, F3) are annotated.

The most striking suppressions of exchange rates of the autoinhibited Pyk2 FERM-kinase manifest at surfaces in the kinase C-lobe and FERM F2 subdomain (Figure 3b). The C-lobe and F2 subdomain surfaces are also the primary interface of the autoinhibited FAK FERM-kinase structure.^*34*^ Indeed, alignment of Pyk2 kinase and FERM domain structures to the FAK FERM-kinase autoinhibited conformation reveals close agreement between the FAK interface and the Pyk2 exchange rate perturbations (Figure 4). The kinase exhibits the largest exchange rate decreases in the G helix (Figure 4, inset 4) and helix EF adjoining the activation loop (Figure 4, inset 5). The perturbations of the FERM F2 subdomain are lower in magnitude and more diffuse (Figure 4, insets 2 and 6). Importantly, HDX-MS is highly sensitive to protein dynamics, and the perturbations may reflect conformational perturbations beyond the direct interface.

**Figure 4.**
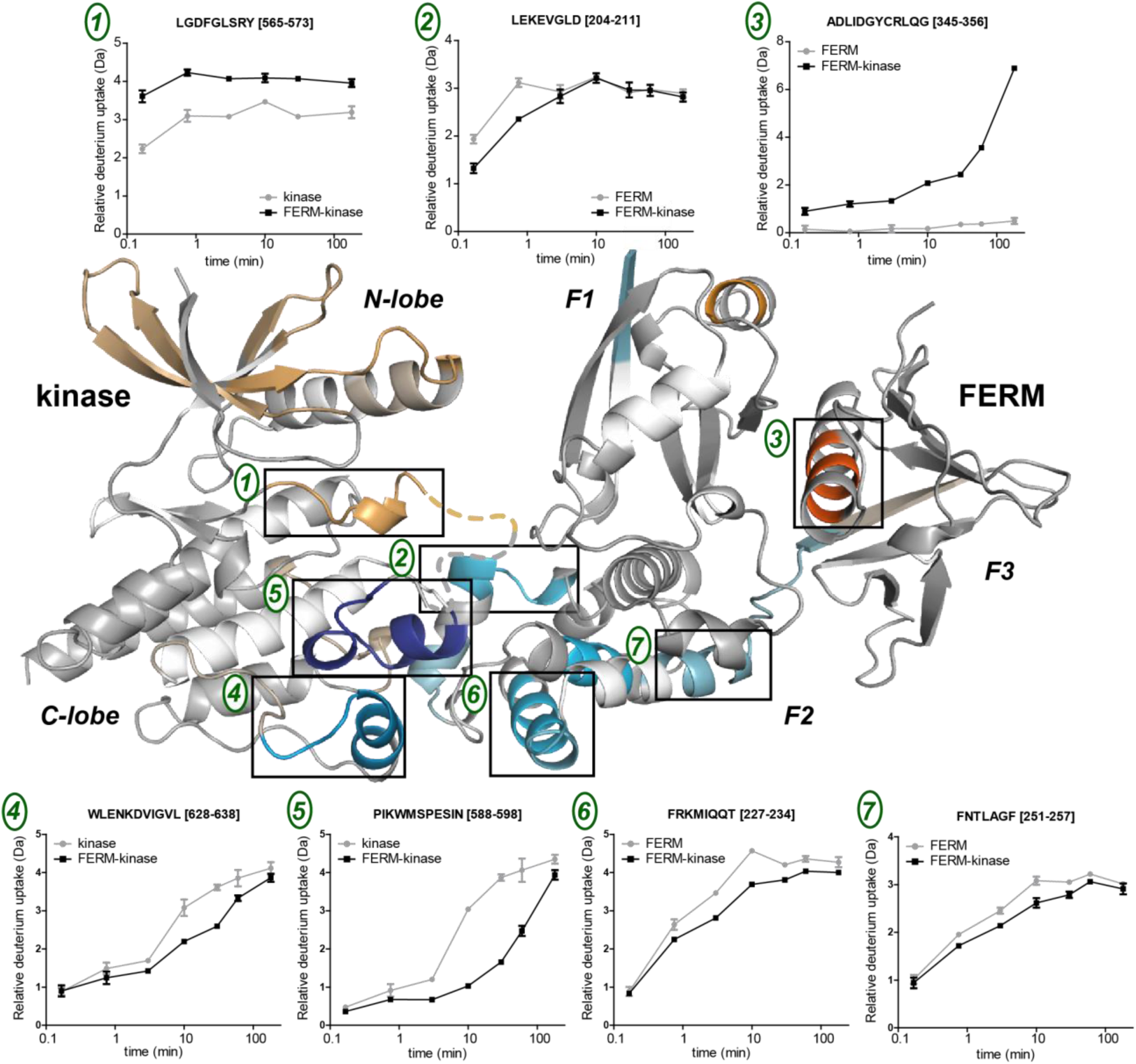
HDX-MS exchange rate perturbations mapped to Pyk2 domains aligned in the autoinhibited conformation of the FAK FERM-kinase (PDB 2j0j). Representative deuterium uptake plots are shown for peptides derived from the featured, numbered regions. Error bars represent standard deviation of three technical replicates.

The autoinhibited conformation also impinges the dynamics of the activation loop (Figure 4, inset 1). The activation loops of the autoinhibited FAK FERM-kinase and isolated Pyk2 kinase are not resolved in crystallography-derived structure. Nevertheless, the exchange rate suppression suggests that the autoinhibited conformation directly perturbs the accessibility or dynamics of the activation loop. Activation loop dynamics are likely to be critical to the regulation of both Pyk2 and FAK. For example, the phosphorylated FAK activation loop adopts a conformation sterically incompatible with the autoinhibited FERM:kinase interface.^*34*^

Notably, the FERM domain perturbations in the autoinhibited conformation overlap and adjoin a basic patch conserved in both Pyk2 and FAK (Figure 4, inset 7). The basic patch is distinct from the FAK FERM:kinase interface, but this region has been implicated in FAK activation^*38*^ via interactions with the c-Met receptor tyrosine kinase^*51*^ membrane phosphoinositides^*32*^, and FAT domain.^*31*^ The impact on the basic patch suggests allosteric communication between the direct FERM:kinase interface and this functionally important surface. Indeed, it is currently unknown how activational stimuli release the autoinhibitory FERM:kinase interaction in either FAK or Pyk2. Allosteric coupling between basic patch and interface may provide a mechanism for activation. On a related note, the core alpha helix of the Pyk2 FERM F2 subdomain has been implicated in Ca^2+^/calmodulin binding and activation of Pyk2.^*35*^ HDX-MS reveals slight perturbations in this helix. However, the solvent accessibility of this helix is minimal, and it is unclear how calmodulin could access this proposed binding site.

Pyk2 FERM and kinase domains exhibit other exchange rate perturbations remote from the putative direct interface. There is faint evidence for perturbations in the FERM F1 subdomain and kinase N-lobe. These regions that make slight contacts with the FERM-kinase linker in the autoinhibited FAK structure.^*34*^ Absence of this linker in the autonomous FERM and kinase constructs may be responsible for the effects in the N-lobe and F1 subdomain. In addition, one anomalous F3 subdomain helix exhibits exchange rate suppression in the autoinhibited conformation (Figure 4 inset 3). Intriguingly, this F3 helix forms an important basic binding cleft for phosphoinositides and membrane protein tails in other FERM-containing proteins.^*52, 53*^ The basic residues are not preserved in FAK and Pyk2, having been substituted with acidic residues. Nevertheless, there may be allosteric linkage between the F3 subdomain and the apparent FERM:kinase interface.

While HDX-MS is exquisitely sensitive to the dynamics of the polypeptide backbone, parallel approaches are helpful for testing the functional importance of conformational perturbations. To validate the autoinhibitory conformation indicated by HDX-MS, we designed a panel of Pyk2 variants with putatively disruptive residue substitutions. We reasoned that disruption of the autoinhibitory conformation would lead to a de-repression of kinase activity.

Eight Pyk2 FERM-kinase variants were generated and purified with residue substitutions engineered at surface sites within and outside the regions identified by HDX-MS (Figure 5A). F187A, E595R, and F599D were selected to probe the C-lobe:F2 subdomain interface observed in our Pyk2 HDX-MS analysis and the FAK FERM-kinase structure.^*34*^ L58A and E404P were selected to test a feature of the FAK FERM-kinase conformation. In FAK, the initial site of autophosphorylation, linker residue Y397, incorporates into a beta sheet in the F1 subdomain. We predicted that substitutions at linker residue (E404P) and F1 beta strand (L58E) could disrupt formation of this possible feature of the autoinhibited conformation in Pyk2. Three residue substitutions (Y265A, L311A, and E342R) were also selected in the F3 subdomain. The F3 subdomain is distant from the autoinhibitory interface in FAK FERM-kinase, yet exchange rate perturbations appeared therein in Pyk2 FERM-kinase.

**Figure 5.**
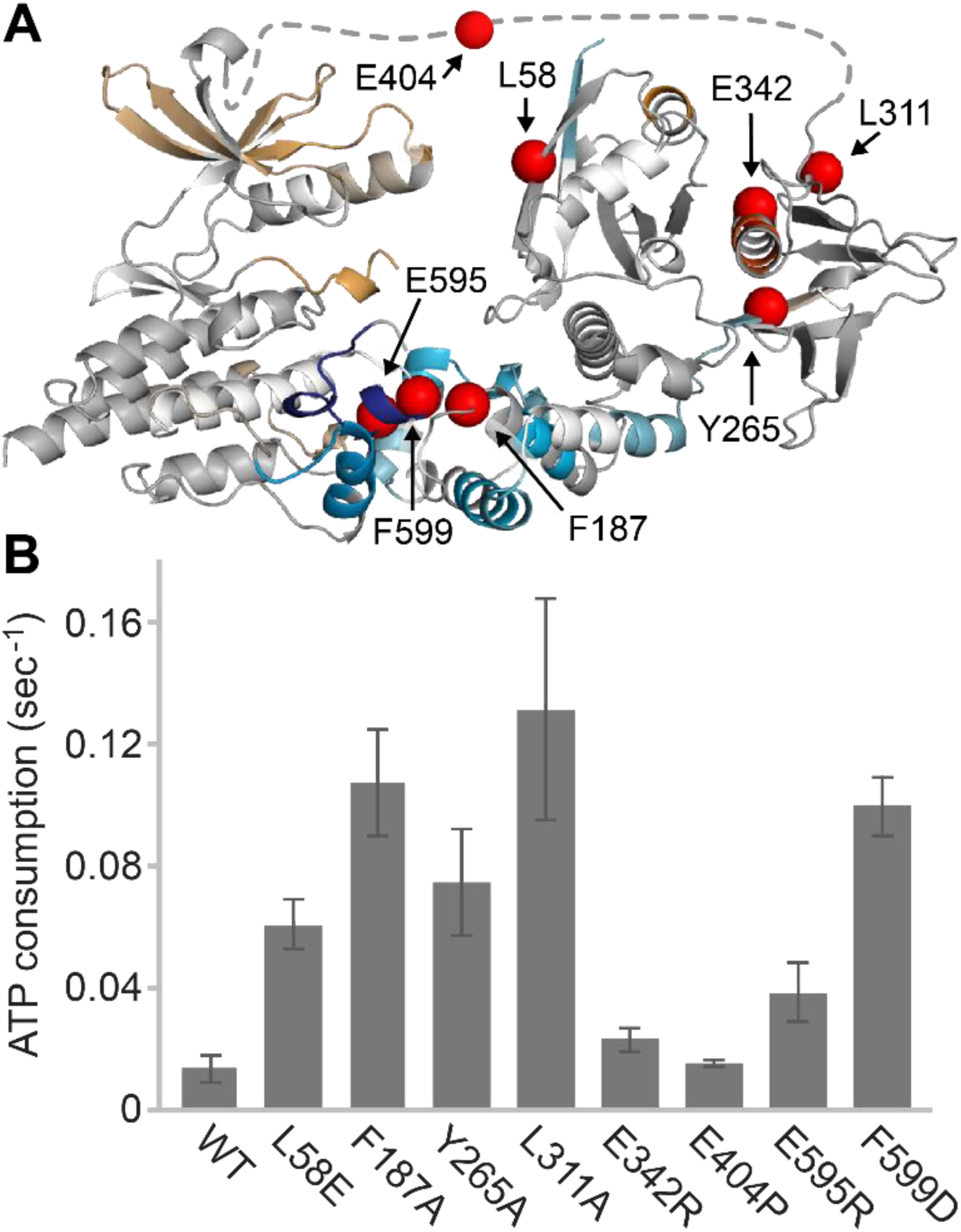
Kinase activity of Pyk2 FERM-kinase variants. (A) Residues selected for putatively disruptive mutations are highlighted as spheres on Pyk2 kinase and FERM domains with HDX-MS exchange rate perturbations mapped. The dotted line represents the FERM—kinase linker. (B) Kinase activity of each purified Pyk2 FERM-kinase variant measured by ATP consumption using Glu:Tyr (4:1) polypeptide substrate at 30 °C. ATP consumption was coupled to NADH oxidation with pyruvate kinase and lactate dehydrogenase in a continuous spectrophotometric assay monitoring absorbance at 340 nm for 5 min. Error bars represent standard deviation of six independent activity assays.

Kinase activity of the Pyk2 variants was measured using an enzyme-coupled, continuous spectrophotometric assay monitoring ADP product formation (Figure 5B). All variants were dephosphorylated with YopH prior to the assay. Basal activity of each variant was assessed using exogenous peptide substrate. Several of the Pyk2 variants exhibited increased kinase activity, indicating a disruption or relaxation of the autoinhibited conformation. F599D and F187A exhibited striking increases in activity, consistent with the importance of the N-lobe:F2 subdomain interface in stabilizing the autoinhibited conformation. Indeed, a F596D FAK FERM-kinase variant, the homologous residue to Pyk2 F599D, exhibited a similar large increase in kinase activity. Nevertheless, E595R exhibited a marginal increase in activity, indicative of some tolerance for variation in the putative interface.

Of the residues designed to test the linker conformation, E404P was indistinguishable from WT basal activity. However, L58E exhibited a nearly five-fold activation. Stable association of the linker with the FERM F1 subdomain main, indeed, play a role in Pyk2 autoinhibition. Alternatively, kinase N-lobe:FERM F1 subdomain contacts may stabilize autoinhibition. However, HDX-MS analysis reports negligible perturbations in this region, and the FAK FERM-kinase structure revealed proximity but little direct contact.^*34*^

Surprisingly, two F3 subdomain variants demonstrated Pyk2 activation. Both Y265A and L311A activated the Pyk2 kinase over five-fold higher than the basal WT activity. E342R was not stimulatory. Interestingly, the strongly activating L311A substitution is adjacent to a Pyk2 residue investigated previously, I308E.^*39*^ In transfected glioma cells, I308E abolishes Pyk2 autophosphorylation. Assuming that the Pyk2 autoinhibitory interface is primarily mediated by F2:C-lobe contacts, the F3 subdomain is likely quite distant from the direct kinase interface. The F3 sensitivity to mutation and kinase-dependent exchange rate perturbation indicates allosteric linkage between F2 and F3 surfaces of the FERM domain.

Taken together, the HDX-MS analysis and activity profiling confirm that FERM domain interaction with the kinase is responsible for Pyk2 autoinhibition. Mapping of the interface suggests an autoinhibitory Pyk2 conformation compatible with the FAK conformation. However, HDX-MS revealed impacts on conformational dynamics distinct from the putative interface, including the functionally important basic patch. In addition, Pyk2 variants designed to disrupt local structure demonstrate that Pyk2 is sensitive to activation by perturbations in all three FERM subdomains. The apparent allosteric communication may confer a mechanism for communicating stimulatory Ca^2+^/calmodulin binding with release of the Pyk2 autoinhibitory interface. It is also possible that allosteric communication can sensitize Pyk2 to regulatory protein interactions and post-translational modifications. We surmise that the allosteric communication between various FERM surfaces and the kinase interface may also have functional relevance to the activation of FAK and other FERM proteins.

## Supporting information

Supporting Information HDX-MS state data

## ACKNOWLEDGMENTS

This work was supported by a grant (award 1715411) from the National Science Foundation, Division of Molecular and Cellular Biosciences. LC-MS/MS instrument acquisition was supported by a gift from the Roy J. Carver Charitable Trust. We thank Graciela Gautier for experimental assistance.

## ABBREVIATIONS

HDX-MS: hydrogen/deuterium exchange mass spectrometry
FERM: band 4.1/ezrin/radixin/moesin

