## Supporting Information S1 and Table 1 for "Conformational dynamics of FERM-mediated autoinhibition in Pyk2 tyrosine kinase"

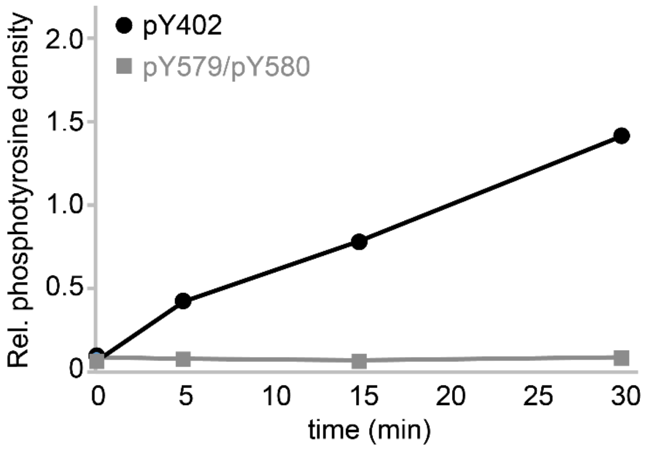


**Figure S1.** Pyk2 FERM-kinase site-specific autophosphorylation. Autophosphorylation was detected by Western blotting using site-specific anti-phosphotyrosine antibodies and quantified by densitometry.

| Data Set | FERM | kinase | FERM-kinase |
|---|---|---|---|
| HDX reaction details | 150 mM KCl, 50 mM HEPES, 2 mM DTT, pD 7.40, 90% $D_2O$, 24 °C | 150 mM KCl, 50 mM HEPES, 2 mM DTT, pD 7.40, 90% $D_2O$, 24 °C | 150 mM KCl, 50 mM HEPES, 2 mM DTT, pD 7.40, 90% $D_2O$, 24 °C |
| HDX time course | 0.167, 0.75, 3, 10, 30, 60, 180 | 0.167, 0.75, 3, 10, 30, 60, 180 | 0.167, 0.75, 3, 10, 30, 60, 180, O?N |
| HDX control samples | Maximum-labeling estimated by 18 hour on-exchange time point (FERM-kinase), n=2 | | |
| Back-exchange (mean / IQR) | 36% / 12% | | |
| # of Peptides | 77 | 52 | 129 |
| Sequence coverage | 92% | 81% | 88% |
| Average peptide length / Redundancy | 10 (2.20) | 11 (2.14) | 10 (1.99) |
| Replicates (biological or technical) | 3 (technical) | 3 (technical) | 3 (technical) |
| Repeatability | 0.083 (average standard deviation) | 0.117 (average standard deviation) | 0.096 (average standard deviation) |
| Significance testing | two-tailed, unpaired t test, $p<0.005$ at time point(s) approximating the middle range of exchange | | |

**Table S1.** HDX-MS Data Summary Table.
